# OrthoPhyl – Streamlining large scale, orthology-based phylogenomic studies of bacteria at broad evolutionary scales

**DOI:** 10.1101/2023.06.27.546815

**Authors:** Earl A Middlebrook, Robab Katani, Jeanne M Fair

## Abstract

There are a staggering number of publicly available bacterial genome sequences (at writing, 2.0 million assemblies in NCBI’s GenBank alone), and the deposition rate continues to increase. This wealth of data begs for phylogenetic analyses to place these sequences within an evolutionary context. A phylogenetic placement not only aids in taxonomic classification, but informs the evolution of novel phenotypes, targets of selection, and horizontal gene transfer. Building trees from multi-gene codon alignments is a laborious task that requires bioinformatic expertise, rigorous curation of orthologs, and heavy computation. Compounding the problem is the lack of tools that can streamline these processes for building trees from large scale genomic data. Here we present OrthoPhyl, which takes bacterial genome assemblies and reconstructs trees from whole genome codon alignments. The analysis pipeline can analyze an arbitrarily large number of input genomes (>1200 tested here) by identifying a diversity spanning subset of assemblies and using these genomes to build gene models to infer orthologs in the full dataset. To illustrate the versatility of OrthoPhyl, we show three use-cases: E. coli/Shigella, Brucella/Ochrobactrum, and the order Rickettsiales. We compare trees generated with OrthoPhyl to trees generated with kSNP3 and GToTree along with published trees using alternative methods. We show that OrthoPhyl trees are consistent with other methods while incorporating more data, allowing for greater numbers of input genomes, and more flexibility of analysis.

**Availability and Implementation:** Code used in this manuscript is available at https://github.com/eamiddlebrook/OrthoPhyl/blob/OrthoPhyl_1.0/. Installation and execution instructions are provided in the associated github README.md file. Third party software versions and OrthoPhyl execution files will remain static in the *OrthoPhyl_1.0* branch, with the *main* branch housing current development. For versions of software dependencies see **Supplemental Table 1**. To aid in usability, a Singularity container is available at https://cloud.sylabs.io/library/earlyevol/default/orthophyl or with the command *singularity pull library://earlyevol/default/orthophyl:1.0_ms*. Also see the GitHub page for Singularity usage guide. Assemblies used within this manuscript are available from https://www.ncbi.nlm.nih.gov/assembly/ with accessions in **Supplemental tables 2-4**.

## 1 Introduction

Underpinning every aspect of an organism’s biology is its evolutionary history. Phylogenetic methods strive to reconstruct these histories using genetic data. With the advent of “Next-” and “Third-gen” sequencing methodologies, the amount of genetic data in public databases has increased dramatically. At writing there are currently 2, 018, 201 bacterial genome assemblies available through NCBI alone (ncbi.nlm.nih.gov/assembly/?term=bacteria). The wealth of available genetic data has transformed the fields of bacterial phylogenetics and taxonomy (Konstantinidis and Tiedje, 2007; Jain *et al*., 2018; Varghese *et al*., 2015).

Unfortunately, most whole genome phylogenetic analysis methods require specialized bioinformatic data processing and expertise (Smith, 2013; Lozano-Fernandez, 2022). There are few very large scale trees to place these sequences into an evolutionary framework. The available large-scale trees are under-resolved and build by reconciling many disparate phylogenetic studies, such as NCBI taxonomy (Schoch *et al*., 2020) or are generated with methods unsuited for narrow and broad evolutionary distances (Hördt *et al*., 2020) reducing their utility at multiple evolutionary scales. This flood of genomic data necessitates an easy-to-use phylogenetic analysis pipeline to help reveal the evolutionary context for the myriad of sequenced bacteria.

### 1.1 Analysis options

#### 1.1.1 Single and Multi-locus

A classic way to generate bacterial trees is to compare single homologous loci (e.g., 16S ribosomal, recA, gyrA, rpoB, or dnaK genes). While largely replaced by other methods, single gene trees have some advantages, which include the ability to use conserved primer sites to amplify sequences, very low computational intensity, and the availability of curated databases (SILVA, RDP, NCBI GenBank, etc.). Using genes other than rDNA allows for greater resolution at certain evolutionary distances but still lacks broad resolution (i.e., Near resolution for swiftly evolving genes, far for conserved) (Yang, 1998). However, these trees are prone to misleading topologies due to horizontal gene transfers, incomplete lineage sorting, paralogous gene conversions, and limited phylogenetic information (Huerta-Cepas *et al*., 2007). Using multi-locus alignments to build phylogenetic trees increases the total amount of phylogenetic information (more sequence used), reduces the effects of incongruent gene tree topologies due to recombination or horizontal gene transfer, and widens the breadth of evolutionary resolution (Gontcharov *et al*., 2004). However, in multi-locus analyses of taxa without well vetted multi-locus methods, locus selection is non-trivial and the effort and cost of primer design and amplification optimization scales with the number of sequences used, which can rapidly become prohibitive for many labs.

Additionally, strains of many taxa, like Brucella, are not differentiated well by available multi-locus methods, leading to unknown evolutionary histories (Sankarasubramanian *et al*., 2019).

#### 1.1.2 Whole genome

Classically, whole genome alignments were required to identify Single Nucleotide Polymorphisms (SNPs) of assemblies (see Shakya *et al*., 2020; Treangen *et al*., 2014 for method details). These methods add considerable phylogenetic data for tree building and can produce different topologies than multi-locus alignments, illustrating their potential utility (Baltrus *et al*., 2014). A major drawback of whole genome alignments is that they tend produce short alignments, or fail outright, for more distantly related genomes (Shakya *et al*., 2020; Chung *et al*., 2018; Darling *et al*., 2004; Angiuoli and Salzberg, 2011). Thus, phylogenetic information drops quickly as a function of evolutionary distance. The practical result is a lack of resolution at even moderate evolutionary distances. For alignments to a reference, having the “missing” data enriched for the more divergent sites will possibly lead to an underestimation of the sequence to reference evolutionary distances for more divergent sequences (Bertels *et al*., 2014; Spencer *et al*., 2007; Shavit Grievink *et al*., 2013). Alternatively, one can use all-vs-all whole genome alignments, however they scale poorly and are limited in the number of input taxa that can be aligned (Angiuoli and Salzberg, 2011).

#### 1.1.3 K-mer overlap

An emergent method to quickly estimate evolutionary histories from genome assemblies is to use *k*-mer content. This involves identifying subsequences of length *k* within each assembly and calculating the shared *k*-mer content (for details see Bussi *et al*., 2021). The advantage of this class of methods is that it is extremely fast relative to alignment-based methods. The comparison of *k*-mer overlap between sequences naturally leads to pairwise distance matrices, thus neighbor-joining trees can be generated directly from the results (Saitou and Nei, 1987). Recent work has integrated more complicated distance statistics with success (Tang *et al*., 2023). Crucially, *k*-mer distances become non-linearly associated with true percent identity, leading to erroneous estimated branch lengths (Jain *et al*., 2018).

An alternative *k*-mer based approach is kSNP (Gardner and Hall, 2013; Gardner *et al*., 2015; Hall and Nisbet, 2023), which identifies SNPs by flanking conserved *k*-mers. This approach allows the use of tree building methods including parsimony, maximum likelihood, and Bayesian. Like whole genome alignments and *k*-mer overlap methods, kSNP suffers from a rapid decline in phylogenetic information as sequences diverge. Additionally, the signal/noise ratio degrades quickly because non-homologous *k*-mers are increasingly common as sequences diverge (Gardner and Hall, 2013).

#### 1.1.4 Predicted gene alignments

To use the wealth of data generated by NGS methods and simultaneously simplify the problem of whole genome alignments, CDS or protein sequences derived from annotations or transcriptome sequencing can be compared. This method breaks the problem of identifying homologous loci into two tractable parts: gene identification, then identification of homologous sequences. This approach identifies many phylogenetically informative sites (like whole genome), but alignments are more tractable (like single loci). However, gene-based alignments are negatively affected by inferred paralogs, i.e., duplicated genes or contamination. Thus, most methods filter out gene families with paralogs. However, see ASTRAL-Pro for a method that is “paralog aware” (Zhang *et al*., 2020). A widely used pipeline that accomplishes this task is GToTree, which annotates assemblies, uses precomputed hidden Marcov models to identify single copy orthologs, aligns predicted amino acid sequences, then generates trees with FastTree2 or IQTree2 from concatenated alignments (Lee, 2019). While focused on protein alignments, GToTree can also be run in nucleotide mode to use CDS alignments for phylogenetic inference instead.

A trade-off arises while using predicted coding or protein sequences to build species trees. Coding sequences (CDS) have the most phylogenetic information due to codon degeneracy but are hard to align at high divergencies (States *et al*., 1991). Proteins remain alignable at high divergences but lack the information content of CDS alignments. This can be addressed by converting protein alignments to corresponding codon alignments, leveraging nucleotide phylogenetic information and protein alignment accuracy (Wernersson and Pedersen, 2003; Bininda-Emonds, 2005). However, at increasing evolutionary distances, even with accurate alignments, mutational saturation in nucleotide data can negatively affect phylogenetic estimates. Evolutionary model selection and dense taxon sampling can largely alleviate these issues, see Kapli *et al*., 2023 for details.

### 1.2 Benefits of an automated workflow

The skills required to generate bacterial phylogenetic trees from whole genomes are extensive. Some of the many steps include gathering and annotating assemblies, identifying and filtering orthologs, sequence alignment, trimming and concatenation, and finally, tree inference (Ashford *et al*., 2020; Lozano-Fernandez, 2022). Each one of these steps, especially for large data sets, requires expertise in picking parameters, file management, and data format manipulation and filtering. Many of these steps also require familiarity with UNIX command-line.

Beyond taxonomic studies, the phylogenetic context of organisms is becoming increasingly important for standard molecular and evolutionary studies. Even with the available data and a clear need for the phylogenetic placement of focal study species, technical barriers preclude many researchers from inferring their evolutionary histories. An easy-to-use, accurate phylogenomic pipeline will encourage wide adoption of phylogenomic methods to complement ongoing environmental, evolutionary, and clinical microbiological research, and perhaps help standardize the estimation of phylogenetic trees within and between research labs.

To meet this demand, we present OrthoPhyl, a phylogenomic pipeline that takes bacterial genomes as input, annotates them, identifies orthologs, converts protein to nucleotide alignments, and builds species trees with both concatenated alignments and gene tree to species tree reconciliation. The workflow accepts an arbitrarily large number of input genomes (tested up to 1200 here). To accelerate analysis, a subset of samples representing the whole dataset’s diversity are identified, their proteomes are used to identify orthologs and build hidden Markov models, which are then expanded to the full input dataset by iterative searches. This strategy allows the generation of trees for 689 Brucella assemblies in ∼48hrs using 30 cpus and 58.3GB memory with no hands-on time required. This pipeline is designed to be an easy to install, scalable, turn-key solution for generating high-resolution bacterial trees from diverse clades.

## 2 Methods

### 2.1 OrthoPhyl’s Workflow

The structure of OrthoPhyl can be broken down into four main steps: Genome assemblies are annotated (Figure 1A), resulting proteins are assigned to orthogroups (Figure 1B), orthogroup proteins are aligned and converted to codon alignments (Figure 1**.C**), and finally, concatenated codon alignments are used to infer phylogenetic trees (Figure 1**.D**). All processes bellow use default parameters unless specified otherwise.

**Figure 1.**
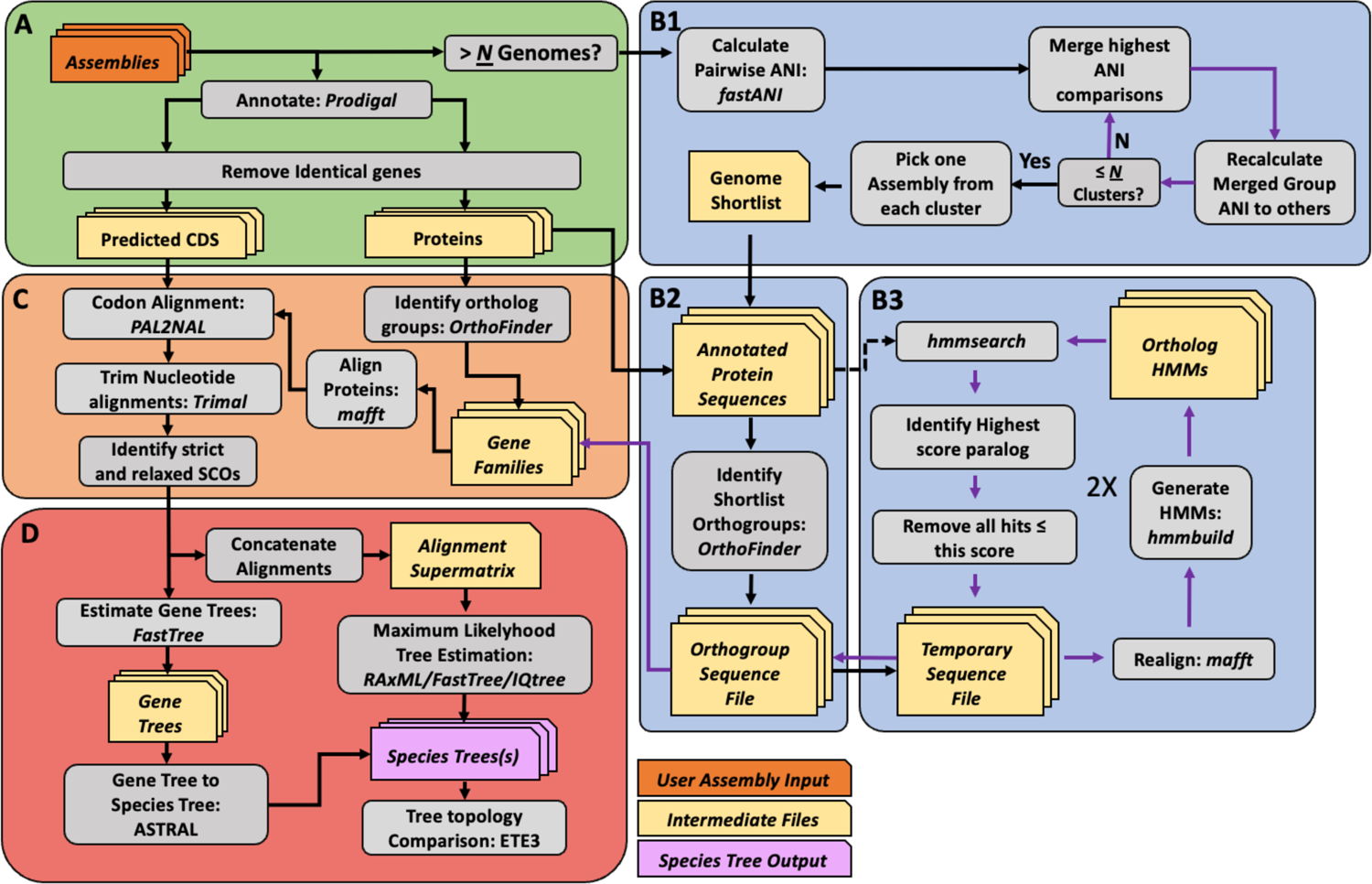
Workflow diagram of OrthoPhyl. Grey boxes indicate processes. Programs used in each step are listed unless a custom script was used. Orange, tan, and purple boxes represent user input, intermediate files, and species tree outputs, respectively. Purple arrows show iterative approaches. The workflow is divided into four main tasks A) annotate assemblies and remove identical CDSs. If more than “N” assemblies are being analyzed, B1) identify a subset of diversity-spanning assemblies, B2) pass them through OrthoFinder to generate orthogroups, and B3) expand the OrthoFinder-identified orthogroups to the full dataset of assemblies through iterative HMM searches. C) Align full orthogroup protein sets, generate and trim matching codon alignments, then filter orthogroups by taxon representation. Finally, D) estimate species tree topologies with concatenated codon alignment supermatrices along with a gene tree to species tree consensus method.

#### 2.1.1 Structural Annotation

OrthoPhyl starts by generating gene calls from input assemblies (Figure 1A) with Prodigal (Hyatt *et al*., 2010). Redundant CDSs (> 99.9% nucleotide identity) found in a genome are removed using bbmap’s *dedupe* (BBMap, 2022) with the logic they are likely the result of recent gene duplications and provide no or little phylogenetic information. This removes unnecessary paralogs from the dataset to preserve the usefulness of the containing orthogroup for downstream analysis.

#### 2.1.2 Orthogroup assignment

To infer orthogroups to use for tree generation, *OrthoFinder* is used with default parameters except for using multiple sequence alignments based gene tree inference (Emms and Kelly, 2015, 2019). When species trees are being generated for large numbers (default: > 30) of genome assemblies, finding orthogroups directly becomes intractable due to the exponential increase in computational time of the all-vs-all homology search. To overcome this, OrthoPhyl identifies a diversity spanning subset of assemblies to identify orthogroups.

Briefly, average nucleotide identity (ANI) is estimated for all assembly pairs with *FastANI* (Jain *et al*., 2018). A custom algorithm is used to cluster assemblies based on pairwise ANI values. Briefly, through successive rounds, samples with the highest ANI are merged into clusters. The merged samples’ ANI to other samples and clusters are averaged. Rounds of merging are performed until N number (default: 30) of clusters are formed. Single representative samples for each of the N groups are chosen as input for OrthoFinder to infer a reduced set of orthogroups. *FastANI* fails to give an ANI percent for genome comparisons with ANI less than ∼75%. For these instances, an arbitrary value of 50% is used. The representative genome picking will be unaffected if there are less than 30 clusters at this or greater distances to all other clusters.

To expand orthogroups to the full assembly dataset, proteins from each orthogroup are realigned with *Mafft* (Katoh and Standley, 2013), then *hmmer* (Eddy, 2008, 2009, 2011) generates hidden Markov models and searches against all predicted proteins using default parameters. OrthoPhyl then finds the minimum HMM hit score cutoff which removes all paralogs, the remaining hits above this score (if any) are designated as the final orthogroup, ensuring all potential paralogs are removed and the greatest number of taxa are represented in the orthogroup. Each orthogroup is then realigned and HMMs are again generated and searched against all proteins to capture additional, more divergent orthologues.

#### 2.1.3 Codon Alignment generation

Once orthogroups are identified for the full set of assemblies, protein sequences are realigned with *Mafft* (Katoh and Standley, 2013). These alignments are then used as the basis for codon alignment using *PAL2NAL* with codon table 11 (Suyama *et al*., 2006). Codon alignments are then trimmed with *trimAL* using arguments “-resoverlap .5-seqoverlap 50-gt .80-cons 60 - w 3” (Capella-Gutiérrez *et al*., 2009). Users can change trimming options with the “control_files.user” if required. *Alignment_Assessment* (Portik *et al*., 2016), is used to visualize the quality of codon alignments, including their phylogenetic signals, taxa per alignment, and alignments per taxa (Figure 2). This is critical for ensuring high-quality data are used for generating trees and can aid in troubleshooting phylogenies with low branch support.

**Figure 2.**
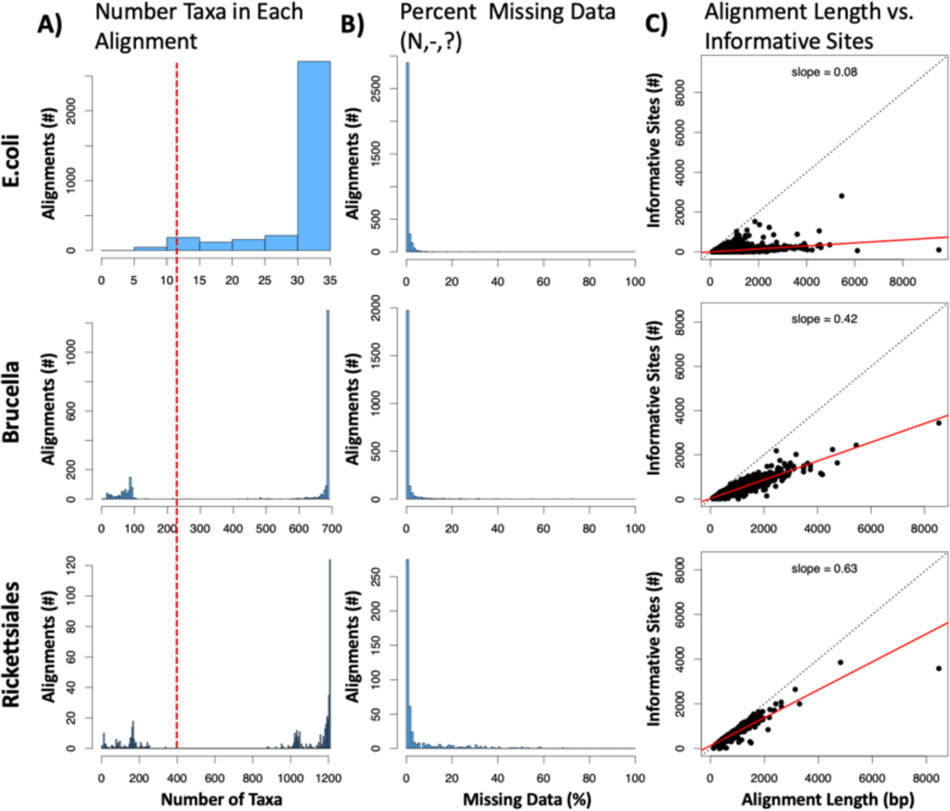
Alignment metric visualization generated by OrthoPhyl. A) Histograms of the number of taxa in each alignment is shown. Dotted red line indicates the >30% taxa cutoff for alignments used to generate the “SCO relaxed” trees. B) Amount of missing data (including gaps and ambiguous bases) per alignment is shown with a historgram. This is exclusive of genomes without detected orthologs. C) The number of phylogenetically informative sites (sites with at least 2 different states in at least 2 taxa each) compared to alignment length for each orthogroup are shown. A linear regression is shown with a solid red line with associated slopes. A one-to-one relationship is shown as a black dotted line.

Our tool then classifies orthogroups as relaxed or strict single-copy orthologs (SCOs). Strict SCOs are genes found in every taxon with no paralogs, while relaxed SCOs are found in a subset of assemblies still with no paralogs. The percent of assemblies with the SCO to count in this set is tunable with the default set as 30% (shown in Figure 2 A: red dashed line). See (Wiens and Morrill, 2011) for analysis on the effects of missing data. These two sets of orthogroups are used in parallel moving forward to generate species trees.

#### 2.1.4 Species Tree Estimation

Two general methods are used to infer species trees for the input assemblies: gene-tree to species-tree estimation with *ASTRAL-III* (Zhang *et al*., 2018) and maximum likelihood methods based on codon alignment supermatrices generated with catfasta2phyml (https://github.com/nylander/catfasta2phyml). First, to use *ASTRAL*, ML gene-trees are generated with either “*FastTree2-gtr-gamma”* (Price *et al*., 2010) or *IQtree2* with default settings (Nguyen *et al*., 2015; Kalyaanamoorthy *et al*., 2017; Hoang *et al*., 2018) per user input. Then ASTRAL will generate species trees with the strict and relaxed SCO gene tree datasets. For the maximum likelihood species tree methods, the user can specify any combination of *IQtree*, *FastTree2 and/*or *RAxML* (Stamatakis, 2014) to infer tree structure(s). Default parameters are used for each tree method except for RAxML and FastTree2 being set to use the GTR + gamma model and 100 and 1000 bootstraps (with <u>Shimodaira-Hasegawa test</u>), respectively, and IQtree2 set to run ModelFinder to identify a best fit evolutionary model and run 1000 ultrafast bootstraps.

Finally, the trees generated during the OrthoPhyl run are compared with generalized Robinson-Foulds metrics provided by ETEtoolkit (Huerta-Cepas *et al*., 2007) indicating the stability of the resultant tree structure when different inference methods and matrix completenesses are used. All default parameters used with these tools can be found in the OrthoPhyl repository at OrthoPhyl/control_file.defaults.

### 2.2 Datasets

As a proof of concept, OrthoPhyl was used to build trees for three bacterial clades. We analyzed well characterized *E. coli*/*Shigella* strains. These included a list of 34 complete genomes analyzed in (Shakya *et al*., 2020), obtained from NCBI.

The genus *Brucella* was used to show OrthoPhyl’s ability to resolve very closely related sequences while dealing with very long relative branch lengths in the same tree (*Ochrobactrum* group). Assemblies were acquired using NCBI Taxon number 234 with the utility script “*gather_genomes.sh*” (Supplemental Methods - Gathering and Filtering Assemblies and **Supplemental** Figure 1) which is packaged with OrthoPhyl (github.com/eamiddlebrook/OrthoPhyl/utils/). This resulted in 689 assemblies after filtering for >98% completion, < 0.1% duplication, and < 1% contamination with CheckM (Parks *et al*., 2015). Here, duplication is the total number of loci identified as marker genes divided by the number of identified marker genes (see Supplemental methods “**Gathering and Filtering Assemblies”** and “utils/checkm_assemblies.slurm” within the github repository). It is important to note that *Ochrobactrum* species were recently moved to the genus *Brucella* (Hördt *et al*., 2020). However, for clarity and due to controversy within the field, we chose to keep the *Ochrobactrum* labeling for these species. An assembly of *Mycoplana dimorpha* (GCA_003046475.1) was added to the dataset as an outgroup.

Finally, a tree for all NCBI Rickettsiales assemblies was generated to illustrate OrthoPhyl’s utility in dealing with bacterial order level divergences. For Rickettsiales assemblies, we again used *gather_genomes.sh,* this time with NCBI taxon number 766. To illustrate a more challenging scenario, filtering of Rickettsiales assemblies was less stringent with cut-offs at > 95% completion, < 0.2% duplication, and < 1.5% contamination, again using CheckM (Parks *et al*., 2015) and our independent calculation of duplication (Supplemental methods “**Gathering and Filtering Assemblies”**. This resulted in 1201 assemblies. Seven *Pelagibacter* assemblies were added to this dataset to serve as an outgroup. Accessions and stats for all genome assemblies used in this manuscript are available in **Supplemental Tables 2-4**.

## 3 Implementation

### 3.1 Small Tree of Closely Related Assemblies: *E. coli and Shigella*

To illustrate Orthophyl’s ability to build trees from moderate numbers of closely related samples on a local Linux computer, we ran our *E. coli* analysis (34 genomes) on a desktop with a 12-core Intel processor with 16 GB RAM running RHEL8 (centOS8). Because the assemblies are of high quality and the stains are closely related, OrthoPhyl was used to generate trees from only the strict SCO dataset. Run statistics are provided in **Table 1**. The full pipeline took 3.25 hours to build trees with FastTree2 and ASTRAL. OrthoPhyl identified 2142 strict SCOs and alignment of these sequences constitutes 2.105 MB with 6% phylogenetically informative sites for the SCO (**Table 1**).

**Table 1.** OrthoPhyl output, runtime, and ram usage.

|  | Assembly Data Set |  |  |
| --- | --- | --- | --- |
| Metric | *E. coli* + *Shigella* | *Brucella* | Rickettsiales |
| Number of assemblies | 34 | 689+1 | 1201+7 |
| Number SCOs - strict | 2217 | 897 | 42 |
| Total Length (bp) | 2,105,000 | 855,924 | 29,248 |
| % Informative SNPs | 6.1 | 41.4 | 81.5 |
| % missing data | 0.4 | 0.5 | 2.3 |
| ANI % (avg/med/min)* | 97.6/99.2/89.9 | 95.4/99.8/63.7 | 83.2/57.3/48.1 |
| Number SCOs – relaxed | - | 1762 | 348 |
| Total Length (bp) | - | 1,627,004 | 295,820 |
| % Informative SNPs | - | 43.3 | 78.9 |
| % missing data | - | 2.7 | 10.6 |
| ANI % (avg/med/min)* | - | 95.0/99.8/63.2 | 82.6/55.7/48.0 |
| CPU time (hr:min) | 16:20 | 373:09:00 | 152:00:00 |
| CPU efficiency | 47% | 24% | 41% |
| Total Runtime (hr:min) | 3:11 | 48:34:00 | 12:10 |
| Max Mem Used (GB) | 13.07 | 58.34 | 17.3 |

\*ANI percentage is calculated with CEANIA (github.com/eamiddlebrook/CEANIA) on the final concatenated codon alignments excluding gapped positions. See Supplemental Methods for details.

The resulting tree built by FastTree2 from the SCO-strict dataset shows successful differentiation of the *E. coli* phylotypes (Figure 3). This tree shows B1 and A as sister phylotypes, with *S. sonnei, S. boydii* and *S. flexneri* between them. That clade is in turn next to phylotype E and *S. dysenteriae*, then D1 and D2. Finally, B2 is sister to all of them. This topology is identical to the reported tree from (Shakya *et al*., 2020) using whole genome alignments, except for the placement of the root (*E. fergusonii)*, where they have the root placed between D2/B2 (red arrow) and the rest while OrthoPhyl’s tree indicates it should be placed between B2 and the other phylotypes.

**Figure 3.**
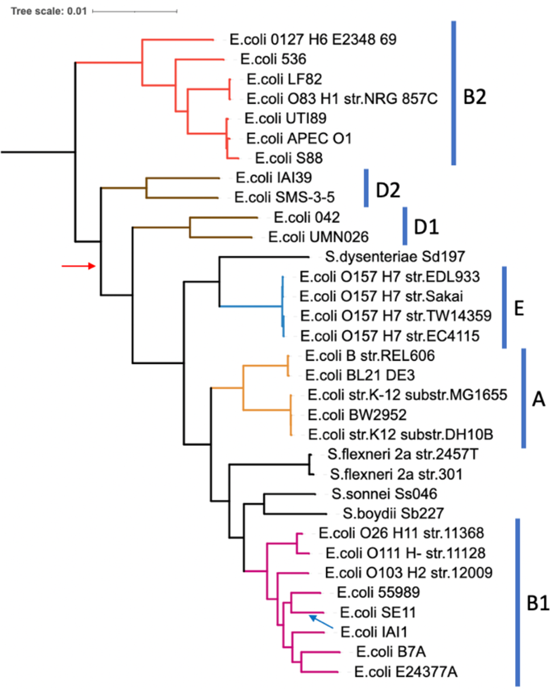
Strict single copy ortholog based maximum likelihood phylogeny of 34 Escherichia and Shigella assemblies inferred by FastTree2 using the GTR gamma model with the default 1000 bootstraps. The tree was constructed with 2217 genes totaling 2.04MB of sequence. Tips are labeled with the given species and strain from NCBI’s BioSample database (details in Supplemental Table 2). All bootstrap support values are 100% except for the split leading to *E. coli* strains APEC O1 and S88 (black arrow), which is 98.7%. Bars next to tree indicate *E.coli* phylotypes [Shakya 2017]. The tree was rooted at *E. fergusonii* then root node was removed. The blue and red arrows indicate the alternative placement of E. coli IAI1 and the root from Shakya 2017, respectively.

### 3.2 Genus level Species trees of 689 assemblies: *Brucella/Ochrobactrum*

For the 689 Brucella/Ochrobactrum assemblies analyzed (plus Mycoplana dimorpha outgroup), OrthoPhyl was run on a RedHat8 compute node with 30 CPUs and 500 GB of ram. The full analysis (generating strict and relaxed SCO trees with *FastTree2* and *ASTRAL*) took just over 2 days (48hrs 31 min). Total ram usage was moderate at 58GB. CPU usage efficiency was 24%, with a total CPU time of approx. 373 hours (**Table 1**).

OrthoPhyl identified 785 strict and 1635 relaxed SCOs in the full dataset of assemblies. The distribution of taxon number represented in each alignment shows that most orthogroup alignments have all taxa represented, with approximately 1400 orthogroups having 680-689 taxa (Figure 2A). The orthogroup alignments have every little missing data (gaps or ambiguous bases) with the vast majority showing less than 10% (Figure 2B). Figure 2C shows the number of informative sites vs. each alignment’s length. A slope of 0.42, indicates the alignments are far from the value that would constitute noisy, erroneous alignments.

The maximum likelihood (ML) *Brucella* phylogeny generated by FastTree2 based on the 1635 relaxed SCOs (Figure 4) shows very close agreement with trees generated by (Ashford *et al*., 2020) for the *Ochrobactrum* clades (see **Supplemental** Figure 2 for high resolution tree with accessions, bootstrap values, and species labeled). For instance, it supports two major *Ochrobactrum* clades, Group A - *O. anthropi, O. lupini, O. tritici. O. pecoris, O. oryzae, O, ciceri, O. intermedium, O. pseudointermedium,* and *O. daejeonense,* and Group B - *O. grignonenense, O. pituitosa(pituitosum), O. quorumnocens, O.rhizosphaerae, O. pseudo grignonense, O. thiophenivorans,* and *O. gallinifaecis* (Figure 4 **orange bars)**. Like Ashford et al., our tree supports *O. endophytica* as basal to the other *Ochrobactrum*. However, our tree shows the clade with *O. haematophila, O. soli* and *O. teleogrylli* as sister to Group B *Ochrobactrum* with >90% support instead of Group A, like in Ashford et al. (Figure 4A and Supplemental Figure 2, orange arrows). A major distinction between our tree topology and the one presented in Ashford et al. is that our tree shows the monophyletic Brucella clade splitting the *Ochrobactrum* clade between *O. endophytica* and all other *Ochrobactrum* with >90% support (**Supplemental** Figure 2**, black arrow**). In contrast, the Brucellae in Ashford et al. split *Ochrobactrum* between Group A and B, as in (Hördt *et al*., 2020). This alternative placement of Brucellae is shown in Figure 4A and **Supplemental** Figure 2 with blue arrow. The difference could be a product of having different outgroups (*Mycoplana ramose, M. dimorpha,* and *Rhizobium etli* for Ashford et al. and Mycoplana dimorpha for OrthoPhyl), and our tree being subject to long branch attraction between *Brucella* and the unbroken long Mycoplana branch. Additionally, our tree indicates that *O. thiophenivorans, O. pituitosa,* and *O. rhizosphaerae* samples are para/polyphyletic with >90% bootstrap support for all associated splits (Figure 4A and **Supplemental** Figure 2**, astrisks**), indicating they were possibly mislabeled when uploaded to NCBI.

**Figure 4.**
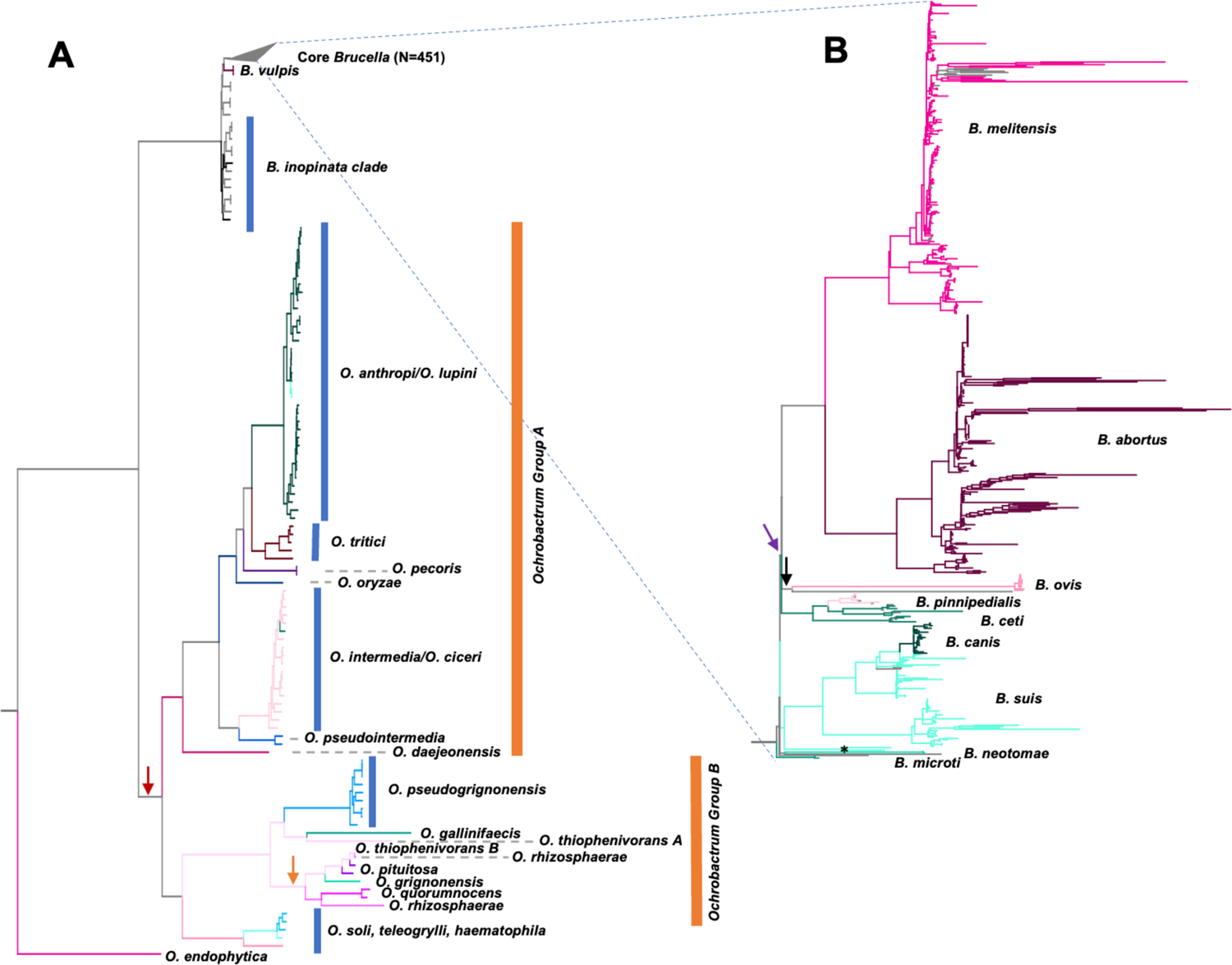
A) Maximum likelihood tree of Brucella/Ochrobactrum assemblies generated by FastTree2 within OrthoPhyl. If provided by NCBI, species are labeled. To aid in visualization of mono/polyphyly, branches are colored by reported species. Internal branch colors are given based on majority rule of offspring. Branch lengths represent relative number of mutations per site. The tree was rooted with *Mycoplana dimorpha,* which was subsequently removed, and the classic Brucella clade was collapsed. B) Subtree of the collapsed classic Brucella clade. A high-resolution version of the full tree showing Species, strain, accession, and bootstrap support is available in Supplemental figure 2.

*Ochrobactrum lupini* and *O. ciceri* are closely associated with *O. anthropi* and *O. intermedia* (respectively). Like Ashford et al. comparing field isolates to type strains, we had many *O. anthropi* and *O. intermedia* assemblies which show that *O. lupini* and *O. ciceri* arose from within their respective clades and are likely misclassified as species, but instead possibly should be classified as *O. anthropi* and *O. intermedia*, respectively. For *O. lupini* and *O. anthropi*, this is supported by additional work (Gazolla Volpiano *et al*., 2019).

The clade comprising the classical *Brucella* agrees with published phylogenies (Figure 4B) (Suárez-Esquivel *et al*., 2020; Ashford *et al*., 2020). *Brucella vulpis* at the base*, B. pinnipedialis* arising from within *B. ceti* (rendering *B. ceti* paraphyletic) (Orsini *et al*., 2022), *B. canis* arising from within *B. suis* (rendering *B. suis* paraphyletic), and *B. melitensis* and *B. abortus* being sister groups. Interestingly, several Brucella assemblies seem to be mislabeled (**Supplemental** Figure 2**, grey tip labels**). Whether these placements are accurate or not remains to be determined, however, B. melitensis and B. abortus are very closely related (precluding saturation) and do not show signs of significant interspecies recombination (Vishnu *et al*., 2015; Suárez-Esquivel *et al*., 2020). The classical *Brucella* have very short branches at the crown (*B. microti*, *B.neotomae*, *B.ovis*, *B. suis*/*canis*, *B. abortus*/*melitensis* and *B. pinnipedialis*/*ceti*), indicating very rapid divergence times for these clades, as seen elsewhere (Ashford *et al*., 2020; Suárez-Esquivel *et al*., 2020; Orsini *et al*., 2022). While the topology of this tree is congruent with several published trees, more analyses are required to create the most robust tree possible (see Discussion).

#### 3.2.1 GToTree vs. OrthoPhyl: *Brucella/Ochrobactrum*

OrthoPhyl compares favorably to the phylogenomics pipeline GToTree (Lee, 2019). We ran GToTree v1.8.4 in nucleotide mode on the same genome dataset as OrthoPhyl allowing the use of 28 cores while searching for orthologs with the Alphaproteobacteria HMM gene set containing 117 models. All other options were left as default. The full analysis took slightly more time than OrthoPhyl, using 50 hours of real time with a total CPU time of 1057 hours. The alignment length produced was 74.9kb, with 51 million total characters and approximately 1 million being missing/ambiguous.

The GToTree topology for deep splits is very similar to the tree generated by OrthoPhyl above. Specifically, all of the species level splits of the Ochrobactrum group are identical between the trees (**Supplemental** Figures 2 and **3**). Additionally, the Brucella clade splits the main Orchrobactrum clade and O. endophytica in both trees. Notable disagreements between the trees include many strain level differences in the Ochrobactrum species subtrees and the basal Brucella. Additionally, the classical Brucella clade shows much lower resolution in the GToTree phylogeny, with many splits supported in less than 50% of bootstrap trees (**Supplemental** Figure 3**, red dots**). These results are perhaps expected due to their relatively recent radiations and GToTree using a highly conserved gene set from Alphaproteobacteria, likely missing Brucella/Ochrobactrum specific phylogenetic signal which could resolve these relationships.

#### 3.2.2 kSNP4 vs. OrthoPhyl: *Brucella/Ochrobactrum*

To compare performance of OrthoPhyl with an alternative assembly-to-tree method, kSNP4 (Gardner *et al*., 2015; Hall and Nisbet, 2023) was run with the same hardware as OrthoPhyl. See Supplemental methods for details. The resulting kSNP4 matrix for the Brucella/Ochrobactrum dataset has 7.15 million SNPs, which is considerably longer than the maximum genome length. Many of these are likely artifacts of the sequences being divergent enough to have mutations within the flanking k-mers used to identify SNPs. Thus, much of the matrix consists of missing data (∼78%) indicating *k-*mers not being shared between assemblies.

FastTree2 was used externally to generate a tree with the *k*-mer based SNP alignment matrix from kSNP4. Like the OrthoPhyl tree, sites found in less than 30% of assemblies were removed up to the point of removing 80% of total sites. This left 1, 569, 787 SNPs (exactly 20% of original SNPs, with ∼13% total missing data). The trimmed SNP alignment file was then used to generate a tree with FastTree2 using the GTR+gamma model and default parameters. For a high-resolution tree with species, strain, bootstap support and assembly accessions labeled, see **Supplemental** Figure 4.

For deep branches outside of the classic Brucella, this tree shows largely the same topology as the tree generated by OrthoPhyl. Some notable exceptions within the Ochrobactrum include: 1) the placement of O. gallinifaecis and one of the *O. thiophenivorans* being sister to *O. pseudogrignonensis* in the OrthoPhyl tree while it is sister to the clade of *O. pseudogrignonensis, O. quorumnocens, O, pituitosa* (among others) in the kSNP4 based tree (Figure 4A**, orange arrow)**, and 2) *O. daejeonensis* is placed one node more basal in the kSNP4 tree (Figure 4A**, red arrow)**. For the *Brucella* clade, there are 2 major differences. The first is the placement of the root, with kSNP4 showing it coming in-between B. ovis and the rest of Brucella (Figure 4B**, black arrow)**. The second is (given the OrthoPhyl root) *B. neotomae* being placed, along with two B. suis samples (Figure 4B**, asterisk)** on the short branch leading to the *B. melitensis, B. abortus, B. ovis, B. pinnipedialis, B. ceti* clade, instead of sister to all *Brucella* internal to *B. microti* (Figure 4B**, purple arrow)**. Compared to kSNP4, OrthoPhyl had favorable runtime and resource usage requirements. Major differences include max memory usage (58 vs 204 GB), max storage (53 vs 480GB) and runtime (48 vs > 216 hours) for OrthoPhyl and kSNP4, respectively.

### 3.3 Order Level Divergence Tree with 1200 samples: Rickettsiales

The order Rickettsiales was chosen as a challenging taxon to test OrthoPhyl’s ability to deal with wide phylogenetic distances and extensive genome reduction. Additionally, this set of assemblies was not filtered stringently, leaving a total of 1208 assemblies (7 outgroup) of varying quality (see Datasets section above). As with the *Brucella* analysis, OrthoPhyl was run on a 30 cpu compute node with 500 GB of available ram. The program took just 12 hours and 10 minutes to complete using a total of 17.3 GB memory. The CPU usage efficiency was 41%, with a total CPU time of approx. 152 hours (**Table 1**).

Even with this challenging genomic dataset, OrthoPhyl identifies > 29, 000 homologous base pairs from strict SCOs and 295, 000 bp from relaxed SCOs (found in 366 or more assemblies). These homologous sequences come from 42 and 348 genes respectively (**Table 1**). The relaxed SCO FastTree2 ML tree reconstructed from the 1201 Rickettsiales assemblies recovers all valid genera as monophyletic (Figure 5A). The sole paraphyletic genus recovered is *Candidatus Jidaibacter*. Other than *Jidaibacter*, the recovered topology of genera is consistent with previously published analyses (Salje, 2021; Schön *et al*., 2022), with two major clades 1) *Wolbachia* sister to *Ehrlichia* and *Anaplasma*, with *Neoricketsia* basal to them and 2) *Rickettsia* and *Orientia* separated from the rest by the root. A high-resolution tree is provided in **Supplemental** Figure 5, which provides species, assembly accession, bootstrap support and number of SCOs per sample.

**Figure 5.**
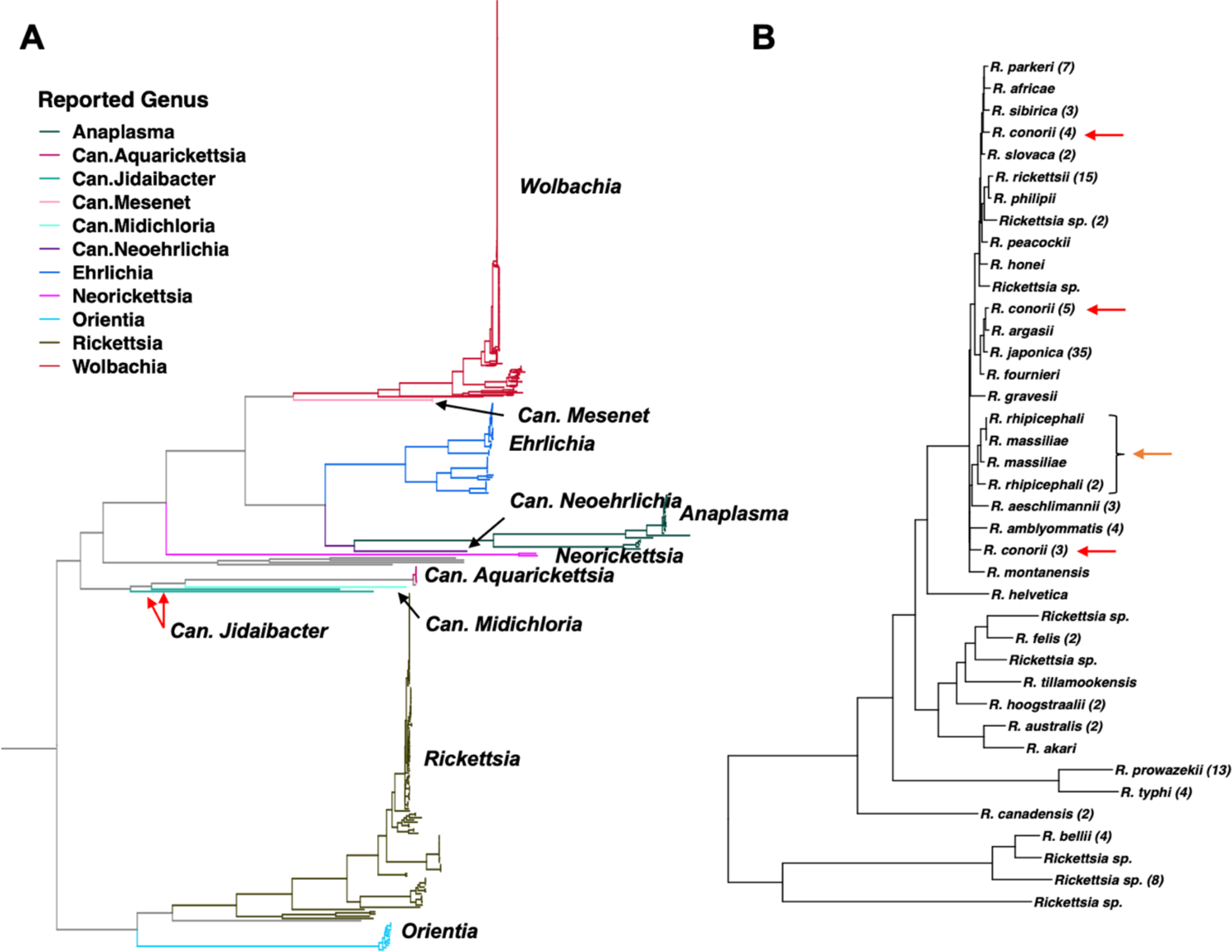
A) Maximum likelihood tree for Rickettsiales generated by FastTree2 within OrthoPhyl. Branch colors show reported genera for each sample according to NCBI metadata. To aid in identification, labels are also provided. Samples with no genus level identification are labeled in grey. Internal branch colors are given based on majority rule of offspring. Note the red Wolbachia clade has been compressed vertically (1/5 ratio) thus clade widths are not proportional to number of samples across the tree. Branch lengths are relative probability of a mutation per site. The tree was rooted with a *Pelagibacter* outgroup that was subsequently removed. Candidate genera (Candidatus) are labeled Can. The single, genus level polytomy (*Canidatus-Jidibacter*) is labeled with red arrows. **B)** The *Rickettsia* subtree is shown with monophyletic species identifiers collapsed. Following species tip labels, in parentheses is the number of collapsed tips. The two most basal branches are omitted, as they are identified only to genus. All assemblies with species level classification show monophyly with the exception of *R. conorii (red arrows)* and *R. massiliae/R. rhipicephali (orange arrow)*.

To illustrate OrthoPhyl’s ability to resolve narrow evolutionary windows along with order level divergences in the same tree, the species rich *Rickettsia* genus subtree is shown (Figure 5B). Only three *Rickettsia* with species level identification (provided by NCBI metadata) show polytomy. First, samples labeled *R. conorii* are placed in 3 very different clades across the tree (Figure 5B**, red arrows)**, indicating that they are possibly mislabeled in NCBI’s genome database. Analysis in Hördt et al, using digital DNA:DNA hybridization, also indicates sequences labeled as *R. conorii* and related species are phylogenenetically problematic. Additionally, the placement of *R. massiliae* and *R. rhipicephali* samples render each other polyphyletic (Figure 5B**, orange arrow**). Other studies mirror our results (Diop *et al*., 2018, 2020). The four R. conorii that are analyzed by Diop et al. are found to be monophyletic in both their papers and this one, with ours having additional R. conorii samples which appear polyphyletic. Diop et al. like us also recovered *R. massiliae* and *R. rhipicephali* as problematic.

It should be noted that, although we recover identical topologies to others, deeper splits in OrthoPhyl’s tree could be affected by saturation at informative sites (**Supplemental** figure 6). Thus, a robust Rickettsiales analysis would involve also inferring trees from alignments excluding saturated genes and/or sites and potentially from corresponding amino acid matrices.

## 4 Key features

### 4.1 Codon alignments

Using codon alignments to infer phylogenetic trees leverages the favorable signal-to-noise ratio of protein alignments (States *et al*., 1991) while gaining phylogenetic signal through codon degeneracy, allowing greater amounts of information per gene (Wernersson and Pedersen, 2003; Bininda-Emonds, 2005; Kapli *et al*., 2023).

### 4.2 Many input assemblies

Whole genome alignment tools, such as Mugsy and Mauve do not scale well for large datasets, with them being limited to generating multiple alignments for about 80 input assemblies (Angiuoli and Salzberg, 2011; Darling *et al*., 2004). OrthoPhyl can generate alignments for >1000 assemblies, allowing large trees to be inferred. It does this by identifying orthogroups in a diversity-spanning subset of assemblies (see Methods), then assigning proteins to these orthogroups for the full set of predicted proteins by iterative HMM searches. This reduces the computationally demanding all-vs-all protein searching that would make these large analyses intractable for many researchers.

### 4.3 User friendly

OrthoPhyl manages the formatting and organization of intermediate files, which is a non-trivial task in a phylogenomic workflow such as this. As with any phylogenetic method, gene sets, alignment and tree estimation parameters will likely need to be tuned to obtain robust results, thus we exposed these variables for altering at runtime. If OrthoPhyl fails due to input error or lack of memory recourses, it can restart at the last completed step. Being robust to restarting also allows users to tune parameters without rerunning the entire pipeline.

## 5 Discussion

OrthoPhyl successfully reconstructs phylogenetic trees for clades with broad evolutionary divergences. This is achieved with little user input; only requiring one command line input pointing the pipeline to the assembly and output directories. OrthoPhyl can generate trees for >30 bacterial assemblies on a laptop or >1000 assemblies on a workstation or single compute node with moderate resources (30 cpus and 100GB RAM). Many useful options are provided by OrthoPhyl, including alignment trimming parameters, number of assemblies from which to identify initial SCOs, tree building software to use, and evolutionary models for tree estimation.

With few exceptions, the trees estimated for the *E. coli/Shigella, Brucella/Ochrobactrum,* and Rickettsiales datasets are consistent with published topologies. This illustrates OrthoPhyl’s ability to resolve broad evolutionary relationships: from highly similar genomes with ANIs of >99% for *E.coli/Shigella* and *Brucella/Ochrobactrum* to highly dissimilar assemblies with ANIs of 57% for the Rickettsiales. With OrthoPhyl being able to generate whole genome-based phylogenies for more than 1000 assemblies, it fills a large gap in phylogenomic analysis software, providing an easy to use, assembly-to-tree tool that will help many researchers incorporate evolutionary analysis into their ongoing bacterial studies.

The *Brucella/Ochrobactrum* and Rickettsiales trees are, to our knowledge, the densest phylogenies published for these clades, let alone using whole genome methods. While more work is necessary to validate the topologies presented here, they are a starting point for assessing metadata-based species labels and expected clade monophyly. Future work will test the robustness of topologies to different evolutionary models, gene sets, tree inference methods, and sample incorporation.

### 5.1 Workflow considerations

There are many parameters to tune during the various steps of phylogenomic inference. Since we focus on usability there is an inevitable tradeoff with optimization, mainly through automated or hard coded parameter choice. OrthoPhyl it does not produce publication ready trees without user inspection of analysis metrics such as alignment quality, single copy ortholog numbers, percent phylogenetically informative sites, mutational saturation and of course branching support values. However, caution should be exercised when interpreting bootstrap support from phylogenomic methods as support values can coalesce on erroneous topologies because of mutational saturation and compositional bias and/or model misspecification. Ultimately, trees should be compared to existing data to ensure major clades are consistent and evidence for novel branching should be closely scrutinized.

Users should take different approaches to inferring trees depending on their final goals. If one wishes to infer an initial phylogenetic tree to inform experimental design or taxon sampling, running OrthoPhyl with default parameters to generate a tree with FastTree2 will likely be sufficient. This will produce a tree using the widely applicable GTR+gamma model. To infer a robust tree for publication, researchers should consider using IQtree2, which runs ModelFinder to choose the evolutionary model which best fits their data. They should also ensure even taxon sampling, with a focus on breaking up long branches. For trees aimed at revising the taxonomy of a studied group, users should consider screening out genes with high saturation and inspecting gene trees for signals of horizontal gene transfer.

Additionally, building trees using multiple “best” models, tree software, and gene sets then comparing results will add robustness to the inferred topologies. We refer readers to (Lozano-Fernandez, 2022) for an in-depth discussion on the subject.

### 5.2 Future

Future versions of this software package will include several additional evolutionary analyses. Bayesian tree inference methods will be incorporated into the pipeline to allow more flexibility in tree estimation. Since codon alignments of single copy orthologs are generated while running this pipeline, we also plan on integrating a software package, e.g., ETE3 (Huerta-Cepas *et al*., 2016), to test evolutionary models such as positive/negative selection and neutrality. A “forest of life” based horizontal gene transfer analysis will be added using gene tree comparisons. This analysis will look at k-means clustering of gene trees to build multiple consensus trees to identify if blocks of genes do not agree with a single consensus species tree hypothesis (Puigbò *et al*., 2019).

In the current iteration of OrthoPhyl, very ancient paralogs (originating before the most recent common ancestor of the dataset species) might be clustered into the same orthogroup, leading to the orthogroup’s removal during filtering even though 2 sub-trees of the orthogroup could be consistent with the species tree. Future iterations of this software will filter the orthogroups in a deep paralog aware manner, using a rapidly generated neighbor-joining species tree to identify and rescue such orthogroups.

Although this pipeline is geared towards prokaryotes, only two changes are needed to adapt it to eukaryotic phylogenetics: 1) allowing users to input precomputed annotations directly, avoiding the complex task of Eukaryotic annotation within the workflow itself and 2) taking into consideration paralog subtrees which are consistent with a consensus species tree. The next version of OrthoPhyl will incorporate these changes to expand the use cases to Eukaryotic phylogenetics.

OrthoPhyl identifies gene family groups for the dataset provided. While this allows the workflow to adapt to any set of genome assemblies, it requires compute resources that could be saved if predefined gene family models could be used as a starting point. Un-clustered protein sequences would then be fed into the standard OrthoPhyl ortholog inference pipeline to capture additional gene families.

### 5.4 Conclusion

Phylogenetic reconstruction is critical to understanding the evolution of pathogenicity, novel traits, horizontal gene transfer, and metabolic potentials of bacteria. Unfortunately, large-scale phylogenomic analyses of diverse bacterial groups require deep understanding of bioinformatic methods and great care in dealing with the myriad intermediate files and formats used during the analysis. To our knowledge, OrthoPhyl is the only software to take >1000 bacterial assemblies with order level divergence, generate ortholog codon alignments, then reconstruct accurate phylogenetic trees without user input or management of intermediate files being necessary. Thus, OrthoPhyl will allow many research groups, including those with modest computing resources and knowledge, to leverage the wealth of publicly available genomic data to enrich their ongoing analyses with robust phylogenomic inferences across a broad swath of bacterial diversity.

## Acknowledgments

The authors would like to thank Migun Shakya, Taehyung Kwon and John Gillece for their valuable insight and comments and the whole Disease Surveillance and Molecular Epidemiology of Brucella project team members in Tanzania and Rwanda, especially Prof. Joram Buza for comments and suggestions. This research was funded by the Defense Threat Reduction Agency through Triad National Security, LLC, operator of the Los Alamos National Laboratory under Contract No. 89233218CNA000001 with the U.S. Department of Energy.

## Bibliography

Angiuoli, S.V. and Salzberg, S.L. (2011) Mugsy: fast multiple alignment of closely related whole genomes. Bioinformatics, 27, 334–342.

Ashford, R.T. et al. (2020) Application of Whole Genome Sequencing and Pan-Family Multi-Locus Sequence Analysis to Characterize Relationships Within the Family Brucellaceae. Front. Microbiol., 11.

Baltrus, D.A. et al. (2014) Incongruence between multi-locus sequence analysis (MLSA) and whole-genome-based phylogenies: Pseudomonas syringae pathovar pisi as a cautionary tale. Mol. Plant Pathol., 15, 461–465.

BBMap (2022) SourceForge.

Bertels, F. et al. (2014) Automated Reconstruction of Whole-Genome Phylogenies from Short-Sequence Reads. Mol. Biol. Evol., 31, 1077–1088.

Bininda-Emonds, O.R. (2005) transAlign: using amino acids to facilitate the multiple alignment of protein-coding DNA sequences. BMC Bioinformatics, 6, 156.

Bussi, Y. et al. (2021) Large-scale k-mer-based analysis of the informational properties of genomes, comparative genomics and taxonomy. PLOS ONE, 16, e0258693.

Capella-Gutiérrez, S. et al. (2009) trimAl: a tool for automated alignment trimming in large-scale phylogenetic analyses. Bioinformatics, 25, 1972–1973.

Chung, M. et al. (2018) Using Core Genome Alignments To Assign Bacterial Species. mSystems, 3, e00236-18.

Darling, A.C.E. et al. (2004) Mauve: Multiple Alignment of Conserved Genomic Sequence With Rearrangements. Genome Res., 14, 1394–1403.

Diop, A. et al. (2020) Genome sequence-based criteria for demarcation and definition of species in the genus Rickettsia. Int. J. Syst. Evol. Microbiol., 70, 1738–1750.

Diop, A. et al. (2018) Rickettsial genomics and the paradigm of genome reduction associated with increased virulence. Microbes Infect., 20, 401–409.

Eddy, S.R. (2009) A new generation of homology search tools based on probabilistic inference. Genome Inform. Int. Conf. Genome Inform., 23, 205–211.

Eddy, S.R. (2008) A Probabilistic Model of Local Sequence Alignment That Simplifies Statistical Significance Estimation. PLOS Comput. Biol., 4, e1000069.

Eddy, S.R. (2011) Accelerated Profile HMM Searches. PLOS Comput. Biol., 7, e1002195.

Emms, D.M. and Kelly, S. (2019) OrthoFinder: phylogenetic orthology inference for comparative genomics. Genome Biol., 20, 238.

Emms, D.M. and Kelly, S. (2015) OrthoFinder: solving fundamental biases in whole genome comparisons dramatically improves orthogroup inference accuracy. Genome Biol., 16, 157.

Gardner, S.N. et al. (2015) kSNP3.0: SNP detection and phylogenetic analysis of genomes without genome alignment or reference genome. Bioinformatics, 31, 2877–2878.

Gardner, S.N. and Hall, B.G. (2013) When Whole-Genome Alignments Just Won’t Work: kSNP v2 Software for Alignment-Free SNP Discovery and Phylogenetics of Hundreds of Microbial Genomes. PLoS ONE, 8, e81760.

Gazolla Volpiano, C., et al. (2019) Reclassification of Ochrobactrum lupini as a later heterotypic synonym of Ochrobactrum anthropi based on whole-genome sequence analysis. Int. J. Syst. Evol. Microbiol., 69, 2312–2314.

Gontcharov, A.A. et al. (2004) Are Combined Analyses Better Than Single Gene Phylogenies? A Case Study Using SSU rDNA and rbcL Sequence Comparisons in the Zygnematophyceae (Streptophyta). Mol. Biol. Evol., 21, 612–624.

Hall, B.G. and Nisbet, J. (2023) Building Phylogenetic Trees From Genome Sequences With kSNP4. Mol. Biol. Evol., 40, msad235.

Hoang, D.T. et al. (2018) UFBoot2: Improving the Ultrafast Bootstrap Approximation. Mol. Biol. Evol., 35, 518–522.

Hördt, A. et al. (2020) Analysis of 1, 000+ Type-Strain Genomes Substantially Improves Taxonomic Classification of Alphaproteobacteria. Front. Microbiol., 11.

Huerta-Cepas, J. et al. (2016) ETE 3: Reconstruction, Analysis, and Visualization of Phylogenomic Data. Mol. Biol. Evol., 33, 1635–1638.

Huerta-Cepas, J. et al. (2007) The human phylome. Genome Biol., 8, R109.

Hyatt, D. et al. (2010) Prodigal: prokaryotic gene recognition and translation initiation site identification. BMC Bioinformatics, 11, 119.

Jain, C. et al. (2018) High throughput ANI analysis of 90K prokaryotic genomes reveals clear species boundaries. Nat. Commun., 9, 5114.

Kalyaanamoorthy, S. et al. (2017) ModelFinder: fast model selection for accurate phylogenetic estimates. Nat. Methods, 14, 587–589.

Kapli, P. et al. (2023) DNA Sequences Are as Useful as Protein Sequences for Inferring Deep Phylogenies. Syst. Biol., 72, 1119–1135.

Katoh, K. and Standley, D.M. (2013) MAFFT Multiple Sequence Alignment Software Version 7: Improvements in Performance and Usability. Mol. Biol. Evol., 30, 772–780.

Konstantinidis, K.T. and Tiedje, J.M. (2007) Prokaryotic taxonomy and phylogeny in the genomic era: advancements and challenges ahead. Curr. Opin. Microbiol., 10, 504–509.

Lee, M.D. (2019) GToTree: a user-friendly workflow for phylogenomics. Bioinformatics, 35, 4162–4164.

Lozano-Fernandez, J. (2022) A Practical Guide to Design and Assess a Phylogenomic Study. Genome Biol. Evol., 14, evac129.

Nguyen, L.-T. et al. (2015) IQ-TREE: A Fast and Effective Stochastic Algorithm for Estimating Maximum-Likelihood Phylogenies. Mol. Biol. Evol., 32, 268–274.

Orsini, M. et al. (2022) Brucella ceti and Brucella pinnipedialis genome characterization unveils genetic features that highlight their zoonotic potential. MicrobiologyOpen, 11, e1329.

Parks, D.H. et al. (2015) CheckM: assessing the quality of microbial genomes recovered from isolates, single cells, and metagenomes. Genome Res., 25, 1043–1055.

Portik, D.M. et al. (2016) An evaluation of transcriptome-based exon capture for frog phylogenomics across multiple scales of divergence (Class: Amphibia, Order: Anura). Mol. Ecol. Resour., 16, 1069–1083.

Price, M.N. et al. (2010) FastTree 2 – Approximately Maximum-Likelihood Trees for Large Alignments. PLOS ONE, 5, e9490.

Puigbò, P. et al. (2019) Genome-Wide Comparative Analysis of Phylogenetic Trees: The Prokaryotic Forest of Life. Methods Mol. Biol. Clifton NJ, 1910, 241–269.

Saitou, N. and Nei, M. (1987) The neighbor-joining method: a new method for reconstructing phylogenetic trees. Mol. Biol. Evol., 4, 406–425.

Salje, J. (2021) Cells within cells: Rickettsiales and the obligate intracellular bacterial lifestyle. Nat. Rev. Microbiol., 19, 375–390.

Sankarasubramanian, J. et al. (2019) Development and evaluation of a core genome multilocus sequence typing (cgMLST) scheme for Brucella spp. Infect. Genet. Evol., 67, 38–43.

Schön, M.E. et al. (2022) The evolutionary origin of host association in the Rickettsiales. Nat. Microbiol., 7, 1189–1199.

Shakya, M. et al. (2020) Standardized phylogenetic and molecular evolutionary analysis applied to species across the microbial tree of life. Sci. Rep., 10, 1723.

Shavit Grievink, L., et al. (2013) Missing Data and Influential Sites: Choice of Sites for Phylogenetic Analysis Can Be As Important As Taxon Sampling and Model Choice. Genome Biol. Evol., 5, 681–687.

Smith, D.R. (2013) The battle for user-friendly bioinformatics. Front. Genet., 4, 187.

Spencer, M. et al. (2007) Conditioned Genome Reconstruction: How to Avoid Choosing the Conditioning Genome. Syst. Biol., 56, 25–43.

Stamatakis, A. (2014) RAxML version 8: a tool for phylogenetic analysis and post-analysis of large phylogenies. Bioinforma. Oxf. Engl., 30, 1312–1313.

States, D.J. et al. (1991) Improved sensitivity of nucleic acid database searches using application-specific scoring matrices. Methods, 3, 66–70.

Suárez-Esquivel, M. et al. (2020) Brucella Genomics: Macro and Micro Evolution. Int. J. Mol. Sci., 21, 7749.

Suyama, M. et al. (2006) PAL2NAL: robust conversion of protein sequence alignments into the corresponding codon alignments. Nucleic Acids Res., 34, W609–W612.

Tang, R. et al. (2023) KINN: An alignment-free accurate phylogeny reconstruction method based on inner distance distributions of *k*-mer pairs in biological sequences. Mol. Phylogenet. Evol., 179, 107662.

Treangen, T.J. et al. (2014) The Harvest suite for rapid core-genome alignment and visualization of thousands of intraspecific microbial genomes.

Varghese, N.J. et al. (2015) Microbial species delineation using whole genome sequences. Nucleic Acids Res., 43, 6761–6771.

Vishnu, U.S. et al. (2015) Identification of Recombination and Positively Selected Genes in Brucella. Indian J. Microbiol., 55, 384–391.

Wernersson, R. and Pedersen, A.G. (2003) RevTrans: multiple alignment of coding DNA from aligned amino acid sequences. Nucleic Acids Res., 31, 3537–3539.

Wiens, J.J. and Morrill, M.C. (2011) Missing Data in Phylogenetic Analysis: Reconciling Results from Simulations and Empirical Data. Syst. Biol., 60, 719–731.

Yang, Z. (1998) On the best evolutionary rate for phylogenetic analysis. Syst. Biol., 47, 125–133.

Zhang, C. et al. (2018) ASTRAL-III: polynomial time species tree reconstruction from partially resolved gene trees. BMC Bioinformatics, 19, 153.

Zhang, C. et al. (2020) ASTRAL-Pro: Quartet-Based Species-Tree Inference despite Paralogy. Mol. Biol. Evol., 37, 3292–3307.

