## Supplementary Figures for "OrthoPhyl – Streamlining large scale, orthology-based phylogenomic studies of bacteria at broad evolutionary scales"

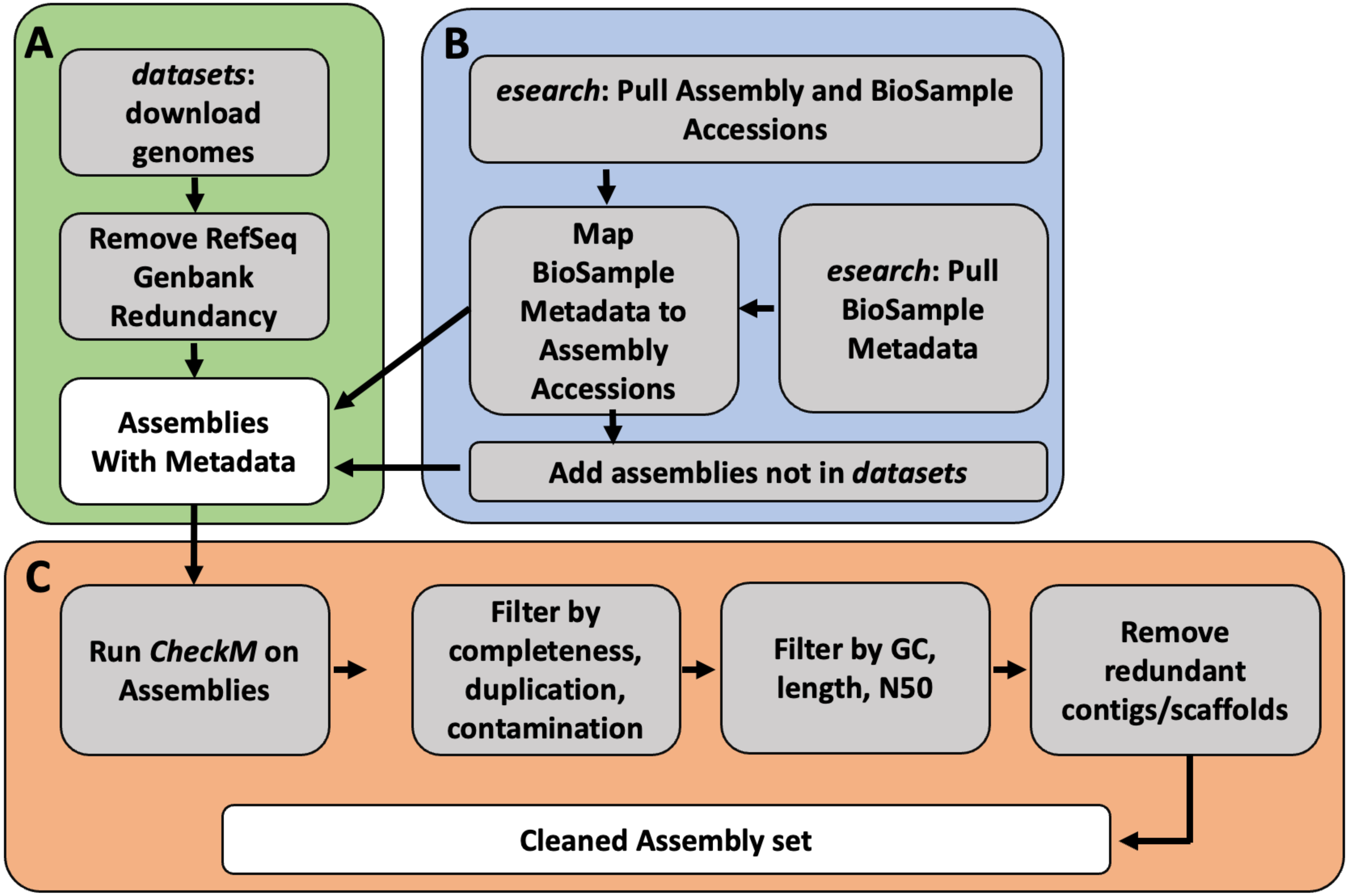


**Supplemental Figure 1**

Workflow diagram of genome gathering and filtering script available from the OrthoPhyl github repository (https://github.com/eamiddlebrook/OrthoPhyl/blob/OrthoPhyl_1.0/utils/gather_filter_asms.sh). The workflow is divided into three tasks **A)** identify and download assemblies, **B)** gather assembly metadata, download assemblies no in NCBI datasets, **C)** calculate genome quality stats and filter to create final set of assemblies to analyze.

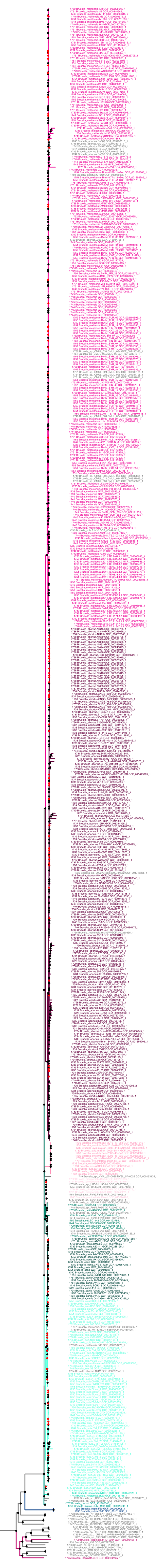

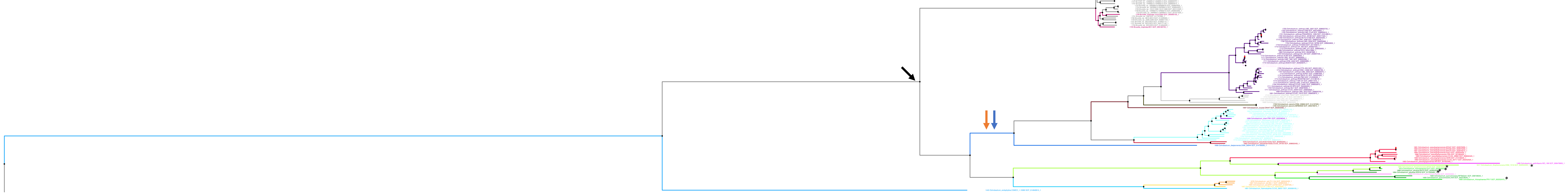


**Supplemental Figure 2–** A high resolution version of main text Figure 3. The tree was estimated for 689 Brucella and Ochrobactrum species using FastTree2 and was rooted with Mycoplana dimorpha (GCF_003046475.1), then leaf was removed. The numbers at tip labels indicate the number of concatenated orthogenes used from each assembly for tree building. Species are labeled by color (to aid in identifying mono- or polyphyly). Points at nodes indicate bootstrap support with black for > 90% and red for < 50% (with unlabeled 90>X>50%). Likely erroneous Brucella species labels have their color removed (i.e. B. abortus clustering with B. melitensis). Finally, accessions are also provided at tips. Orange arrow denotes alternative placements from Ashford et al. of O. haematophila, O. soli and O. teleogrylli, blue arrow shows the alternative placement of the Brucelae from Ashford et al. and Hördt et al.. Black arrow highlights the Brucelae node having >90% bootstrap support.

**Reported Species**

- Brucella_abortus
- Brucella_canis
- Brucella_ceti
- Brucella_inopinata
- Brucella_melitensis
- Brucella_microti
- Brucella_neotomae
- Brucella_ovis
- Brucella_pinnipedialis
- Brucella_suis
- Brucella_vulpis
- Ochrobactrum_anthropi
- Ochrobactrum_ciceri
- Ochrobactrum_daejeonensis
- Ochrobactrum_endophytica
- Ochrobactrum_gallinifaecis
- Ochrobactrum_grignonensis
- Ochrobactrum_haematophila
- Ochrobactrum_intermedia
- Ochrobactrum_lupini
- Ochrobactrum_oryzae
- Ochrobactrum_pecoris
- Ochrobactrum_pituitosa
- Ochrobactrum_pseudintermedia
- Ochrobactrum_pseudogrignonensis
- Ochrobactrum_quorumnocens
- Ochrobactrum_rhizosphaerae
- Ochrobactrum_soli
- Ochrobactrum_teleogrylli
- Ochrobactrum_thiophenivorans
- Ochrobactrum_tritici

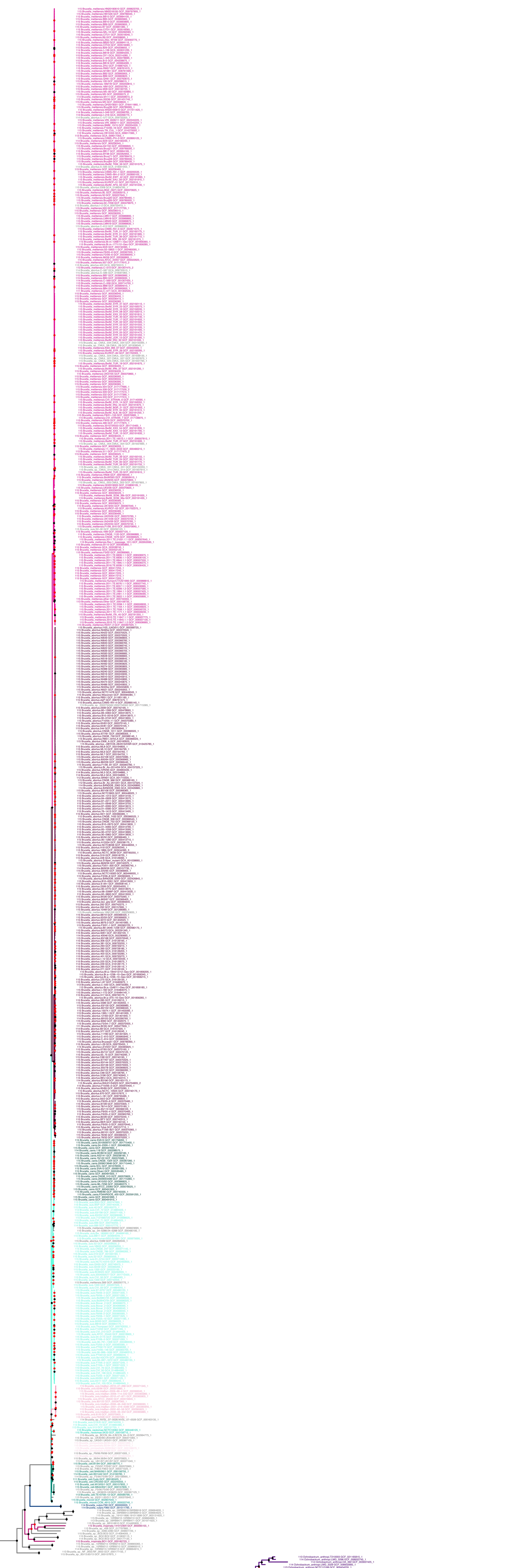

**Supplemental Figure 3 –** The phylogenetic tree estimated for 689 Brucella and Ochrobactrum species using GToTree in nucleotide mode with FastTree2 using the GTR+gamma model and 1000 bootstrap replicates. The resulting tree was rooted with Mycoplana dimorpha (GCF_003046475.1), then leaf was removed. The numbers at tip labels indicate the number of concatenated orthogenes used from each assembly for tree building. Species are labeled by color (to aid in identifying mono- or polyphyly). Points at nodes indicate bootstrap support with black for > 90% and red for < 50% (with unlabeled 90>X>50%). Likely erroneous Brucella species labels have their color removed (i.e. B. abortus clustering with B. melitensis). Finally, accessions are also provided at tips.

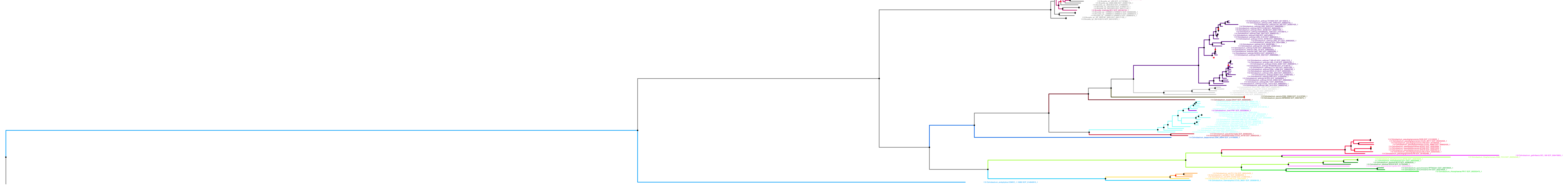

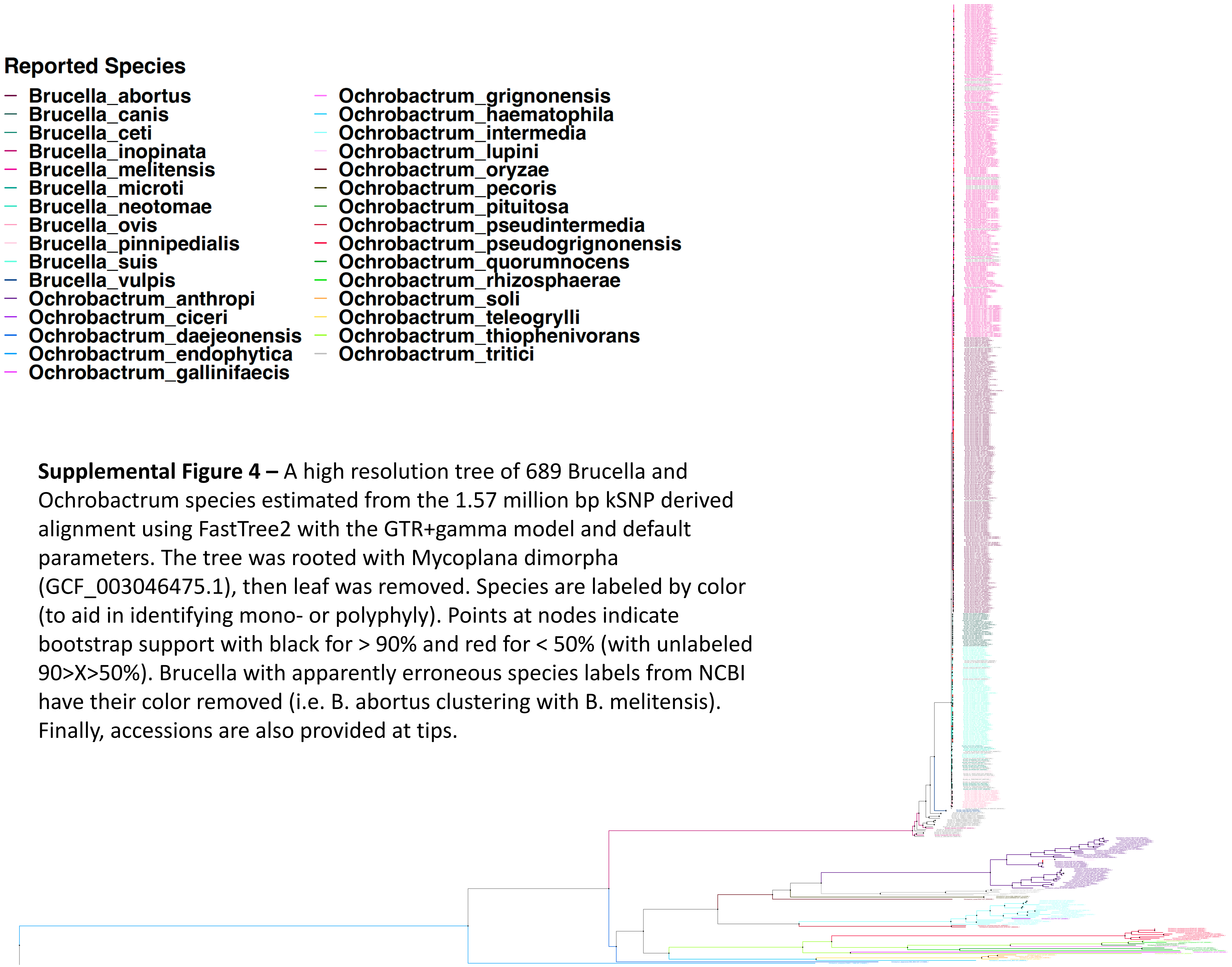


**Supplemental Figure 4 –** A high resolution tree of 689 Brucella and Ochrobactrum species estimated from the 1.57 million bp kSNP derived alignment using FastTree2 with the GTR+gamma model and default parameters. The tree was rooted with Mycoplana dimorpha (GCF_003046475.1), then leaf was removed. Species are labeled by color (to aid in identifying mono- or polyphyly). Points at nodes indicate bootstrap support with black for > 90% and red for < 50% (with unlabeled 90>X>50%). Brucella with apparently erroneous species labels from NCBI have their color removed (i.e. B. abortus clustering with B. melitensis). Finally, accessions are also provided at tips.

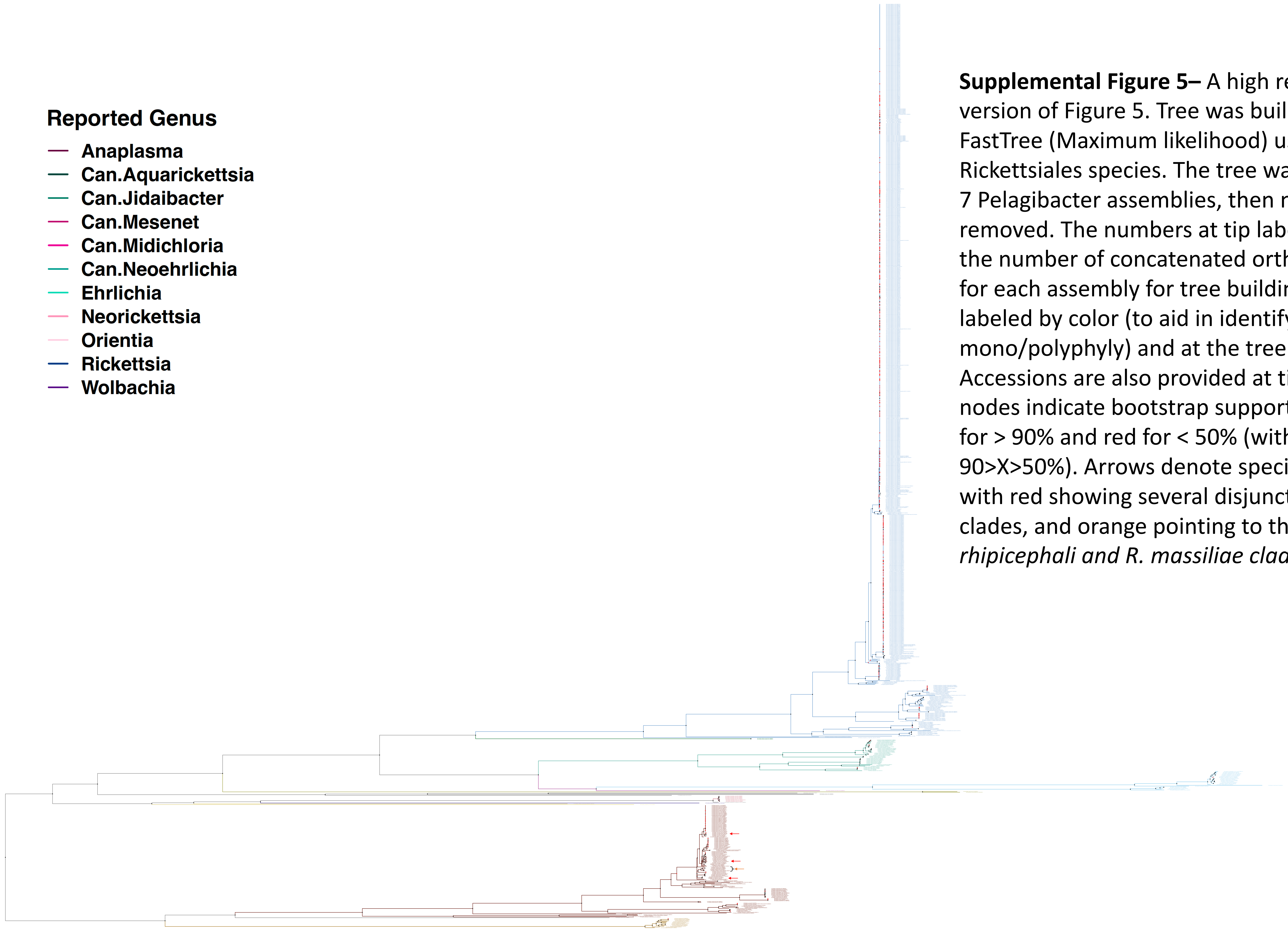


**Supplemental Figure 5–** A high resolution version of Figure 5. Tree was build with FastTree (Maximum likelihood) using 1201 Rickettsiales species. The tree was rooted with 7 Pelagibacter assemblies, then node was removed. The numbers at tip labels indicate the number of concatenated orthogenes used for each assembly for tree building. Species are labeled by color (to aid in identifying mono/polyphyly) and at the tree leaves. Accessions are also provided at tips. Points at nodes indicate bootstrap support with black for > 90% and red for < 50% (with unlabeled 90>X>50%). Arrows denote species polyphyly; with red showing several disjunct R. conorii clades, and orange pointing to the mixed R. *rhipicephali and R. massiliae clade.*

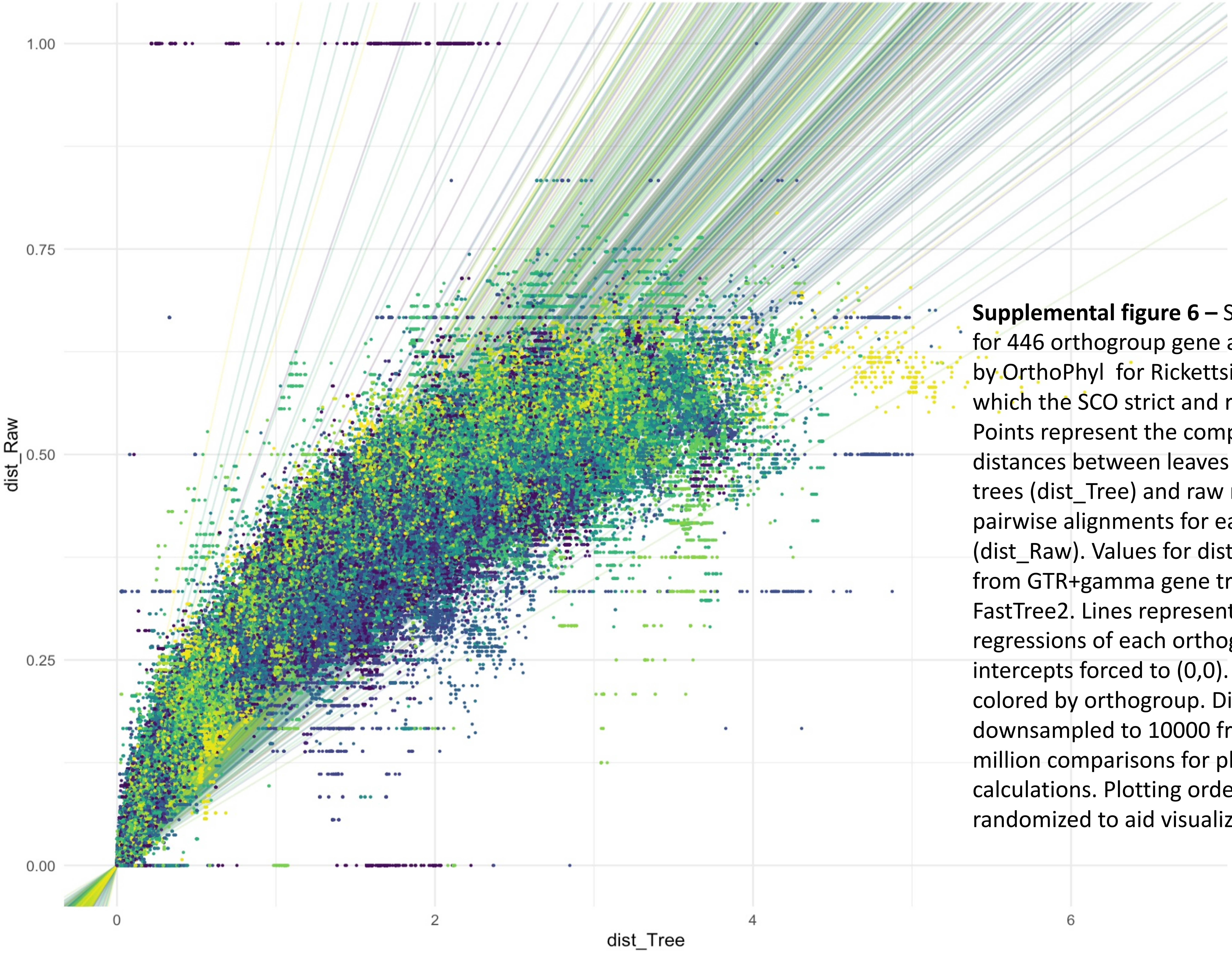


**Supplemental figure 6 –** Saturation estimation for 446 orthogroup gene alignments produced by OrthoPhyl for Rickettsiales assemblies (of which the SCO strict and relaxed set are drawn). Points represent the comparison of phylogenetic distances between leaves on the inferred gene trees (dist_Tree) and raw nucleotide distances of pairwise alignments for each orthogroup's CDSs (dist_Raw). Values for dist_Tree were calculated from GTR+gamma gene trees produced by FastTree2. Lines represent the slopes of linear regressions of each orthogroup with their intercepts forced to (0,0). Points and lines are colored by orthogroup. Distances were downsampled to 10000 from a max of 1.4 million comparisons for plotting and regression calculations. Plotting order of points are randomized to aid visualization of data spread.
