## Supplementary Methods for "OrthoPhyl – Streamlining large scale, orthology-based phylogenomic studies of bacteria at broad evolutionary scales"

**1 Supplemental Methods**

**1.1 Gathering and Filtering Assemblies**

A major hurdle for generating trees with hundreds of assemblies is streamlining the gathering and filtering of sequences. To help users acquire genomes, we have designed a utility script (https://github.com/eamiddlebrook/OrthoPhyl/blob/OrthoPhyl_1.0/utils/gather_filter_asms.sh” that pulls all assemblies from NCBI with the "*datasets*" tool that match a user defined NCBI TaxID **(Supplementary Figure 1A)**. The *datasets* tool leads to redundant RefSeq and GenBank genomes. Therefore, the script automatically removes this redundancy, keeping only the RefSeq entry if both are present.

The utility script also helps users to concurrently gathering assembly metadata with bulk sequence downloads. Since a phylogenetic tree’s scientific utility largely hinges on sample associated metadata, we added modules to pull many useful metadata fields from NCBI (**Supplementary Figure 1B**). In some cases, the required sample information is disassociated from the assembly accession, but available through the BioSample accession. Therefore, the script maps metadata from BioSample accessions to assembly accessions. Not all assemblies from NCBI are available through the *datasets* tool. If assembly accessions are found with *esearch* but are not downloaded with *datasets*, the script attempts to download these assemblies separately.

When building phylogenetic trees, it is critical to use high-quality genome assemblies. This script leverages CheckM (Parks *et al.*, 2015) to gather information on genome completeness and duplication, along with standard assembly stats (scaffold/contig n50, total length, % gaps etc.) (**Supplementary Figure 1C)**. We also added an independent metric of duplication that is (# loci identified as marker gene/number of marker genes). This metric is calculated directly from the CheckM summary statistics, where numbers of marker genes with 1-5+ duplications are listed. It should be noted that, for assemblies with marker genes showing greater than 5 members, this measure will be an underestimate. Users can choose custom filtering parameters based on these outputs. We recommend that users experiment with filtering parameters before settling on any. For instance, filtering on assembly length can be counterproductive for order level divergences where large differences in assembly length are not necessarily due to assembly error (e.g. Spirochettales min length < 1MB, max length > 3.7MB). Additionally, each assembly is screened for contigs that are completely contained within other contigs and fully contained contigs are removed. With the assumption that these are either assembly errors (false duplicates) or that they represent recent sequence duplications, creating 100% identical paralogs, they will not contribute phylogenetic information. Lastly, assemblies that do not pass GenBank and RefSeq databases "taxonomy check" can be filtered out to remove dubious assemblies. However, this feature should be used with caution as some clades always show “Inconclusive”, no matter the assembly quality (e.g. Wolbachia).

**1.3 kSNP**

First, the optimal k-mer length for the dataset was identified with Kchooser (packaged with kSNP4) as length 13. kSNP3 (Gardner *et al.*, 2015) was run using 28 threads and *k*-mer length of 13 with all other paramerters as default. After 9 days of run-time, the analysis crashed with a segmentation fault during the FastTree2 (Price *et al.*, 2010) stage. The excessive runtime was likely a product of the parsimony tree generator, Parsimonitor (github.com/stamatak/Parsimonator-1.0.2), being run in single thread mode, and the segmentation fault could have been a lack of sufficient memory, however, server logs indicate that only 205 of 500 available GB were used. Since the inbuild tree generation was not successful, we used the SNP file to estimate a tree outside of kSNP. With TRIMAL, we trimmed sites that have more than 70% missing data, keeping a minimum of 20% of sites. Because of the high amount of missing data, this left exactly 20% of the data; an alignment of 1.569KB. This trimmed SNP file was then used to build a tree within FastTree2 (Price *et al.*, 2010) using the GTRGAMMA model (**Supplementary Figure 3**).

**1.3 Calculating Exact Average Nucleotide Identity from Alignments (CEANIA)**

To generate the exact ANI values for **Table 1** we developed a python program that calculates all pairwise ANIs for an input multifasta. *CEANIA* is written in python 3 and runs with base python. The software is freely available at github.com/eamiddlebrook/CEANIA. The program works for single gene (short) alignments all the way up to whole genome alignments for bacteria. It can handle an arbitrary number of input sequences. It is multithreaded to decrease total runtime. For this manuscript, we opted to report an ANI which disregards any gaps or ambiguous bases (N,-,?) to have a conservative metric of relatedness.
